# A toxic ankyrin cysteine protease effector RipBH of brown rot triggered autophagy-associated cell death

**DOI:** 10.1101/2023.03.08.531657

**Authors:** Xueao Zheng, Mengshu Huang, Xiaodan Tan, Bingsen Wang, Yanping Li, Hao Xue, Dong Cheng, Huishan Qiu, Wenhao Li, Botao Song, Huilan Chen

## Abstract

Potato brown rot, caused by *Ralstonia solanacearum*, is one of the most destructive diseases of potatoes. The pathogen could hide in the tuber, leading to the rotting tubers. However, few mechanisms of pathogenesis in tubers caused by brown rot were reported. Here, we identified a highly virulent type III effector RipBH, which is not only required for the pathogenesis of potato brown rot but also displays strong cell toxicity in yeast and tobacco. We found RipBH is a novel structural cysteine protease with a large ankyrin repeat domain that contains 10 ankyrin repeats, we named it as an ankyrin cysteine protease. Biochemical analysis showed that all the ankyrin repeats are required for virulence, and the first five ankyrin repeats are indispensable for auto-cleavage site recognition. Further analysis showed that RipBH triggered autophagy-associated cell death. The ankyrin cysteine protease effector existed extensively in plant and animal pathogens suggesting the ankyrin cysteine protease effectors are functionally essential for pathogen pathogenesis. Our study enhances our understanding of this type of cysteine protease and illustrates the pathogenesis of cysteine protease in potato brown rot.

## MAINTEXT

*Ralstonia solanacearum* is a soil-borne bacterium causing the widespread disease known as bacterial wilt. Due to its robust adaptation and survival ability, *R. solanacearum* could infect more than 250 plant species (Genin and Denny, 2012). As significant genetic diversity exists within the *R. solanacearum* species, *R. solanacearum* is also termed as the *R. solanacearum* species complex (RSSC) (Fegan and Prior, 2005; Genin and Denny, 2012). RSSC causes severe economic losses worldwide. The pathogen infects economically important crops such as potato, tomato, eggplant, peanut, pepper, and banana. Potato is the third most important food crop, and is also one of the favorite hosts of RSSC (Lowe-Power et al., 2016; Wang et al., 2017). The major RSSC pathovar infecting potato belongs to phylotype IIB(Van Elsas et al., 2000). This type pathogen has adapted to potato growth environment and causes brown rot of potato tubers (Cellier and Prior, 2010). RSSC enters root through wound, and colonizes in the xylem. It could hide in tuber causing wider spread (Wang et al., 2017). RSSC replicates fast in tuber and causes cell death and necrosis eventually resulting in the rotting tubers (Cellier and Prior, 2010). However, few mechanisms of pathogenesis in tubers caused by brown rot were reported.

One of the *R. solanacearum* powerful weapons for infection is the type III secretion system, which injects type III effectors (T3Es) into the host cytoplasm (Deslandes and Genin, 2014; Peeters et al., 2013). The pan-effectome of the RSSC has been exhaustively identified and is composed of more than 100 different T3Es (Landry et al., 2020; Peeters et al., 2013). To benefit proliferation, pathogens use T3Es to manipulate the host immune response, modulate the host metabolism, and trigger cell death (Feng and Zhou, 2012). Thus, RSSC T3Es provide a large and varied molecular library to understand plant biological process (Cui et al., 2015; Feng and Zhou, 2012). Although T3Es damage cellular processes through multiple mechanisms, the most efficient way to subvert the host cellular environment is using enzyme (Shao, 2008). Cysteine proteases, also named thiol proteases, are hydrolase enzymes that degrade proteins. These proteases share a common catalytic mechanism that involves a nucleophilic cysteine thiol in a catalytic center (Chapman et al., 1997). Cysteine proteases are involved in various biological processes such as growth, development, aging, death, and immunity (Chapman et al., 1997). In bacteria, cysteine enzyme is also used as a powerful weapon to destroy host proteome homeostasis (Shao et al., 2002; Shao et al., 2003).

The first reported bacterial effector protein cysteine protease is YopT in Yersinia (Shao et al., 2002). YopT cleaves Rho GTPases protein, abolishing GTPases anchoring to the membrane, which leads to the disruption of actin cytoskeleton in host cells. The YopT protease family includes other members such as AvrPphB in the *Pseudomonas syringe* (Shao et al., 2003). AvrPphB cleaves the PBS1 protein family, including well-studied receptors like kinase BAK1 (Boller and He, 2009; Yuan et al., 2021). All these effectors belong to papain-like cysteine proteases, which contain prodomain and mature domain (Otto and Schirmeister, 1997). Cysteine protease prodomain usually acts as an endogenous inhibitor of the mature enzyme. To activate the mature enzyme, removal of the prodomain is always necessary (Lecaille et al., 2002). They are usually 23-30 kDa in size, binding the substrate through protein surface interaction (Chapman et al., 1997; Zhu et al., 2004).

Cell death triggered by effectors has been widely reported in plant pathogens. Many effectors triggered cell death through the effector-triggered immunity (ETI) pathways (Zhang et al., 2020; Zheng et al., 2017). For example, the hypertensive response, a type of programmed cell death is triggered by avirulence factors to constrain bacteria growth in the leaves (Lapin et al., 2020). However, the environment of the root is much more complicated than that of leaves, for example, the moisture in the soil is often higher than that in the air and the microbiome composition in the soil is much more sophisticate than in the leaf (Bai et al., 2015). Thus, it’s worthy to know whether the cell death in the roots or tubers is able to constrain the soil-born pathogen replication.

RipBH is a conserved gene in sequenced strains in the phylotype IIB clade (Zheng et al., 2019). To test whether RipBH is a T3E, we used the cyaA report system to analyze the secretion of RipBH of T3SS. The results showed the secretion signal could be detected in the wilt type strain UW551, but not in T3SS mutant strain Δ*hrp*, which suggested RipBH is indeed a T3E of RSSC (Figure S1). To test whether the RipBH is required for the virulence to potatoes, we generate a RipBH mutant strain Δ*RipBH* and the genetic complementation strain Δ*RipBH/RipBH*. Pathogen infection assay showed Δ*RipBH* was compromised in virulence on both potato plants and tubers. In contrast, as shown in Figures 1a and 1b, the disease phenotype of the complementation strain (Δ*RipBH/RipBH*) was similar to that of the wild-type (UW551). These data strongly suggested that RipBH contributes to the virulence of UW551 during the invasion to potato plant. In addition, the morphological analysis showed the tuber cells around inoculation site manifested extensive cell death, and the all of the starch granules were almost destroyed by UW551,compared with Δ*hrp* (Figure 1c-e). Notably, the area of cell death triggered by Δ*RipBH* was significantly smaller than that of wild type UW551. In contrast, the area of cell death triggered by Δ*RipBH/RipBH* was similar to that of the wild type, and the starch granules in these cells were completely degraded (Figure 1c-1e). These data strongly suggested that RipBH promotes tuber cell death caused by UW551.

**Figure 1.**
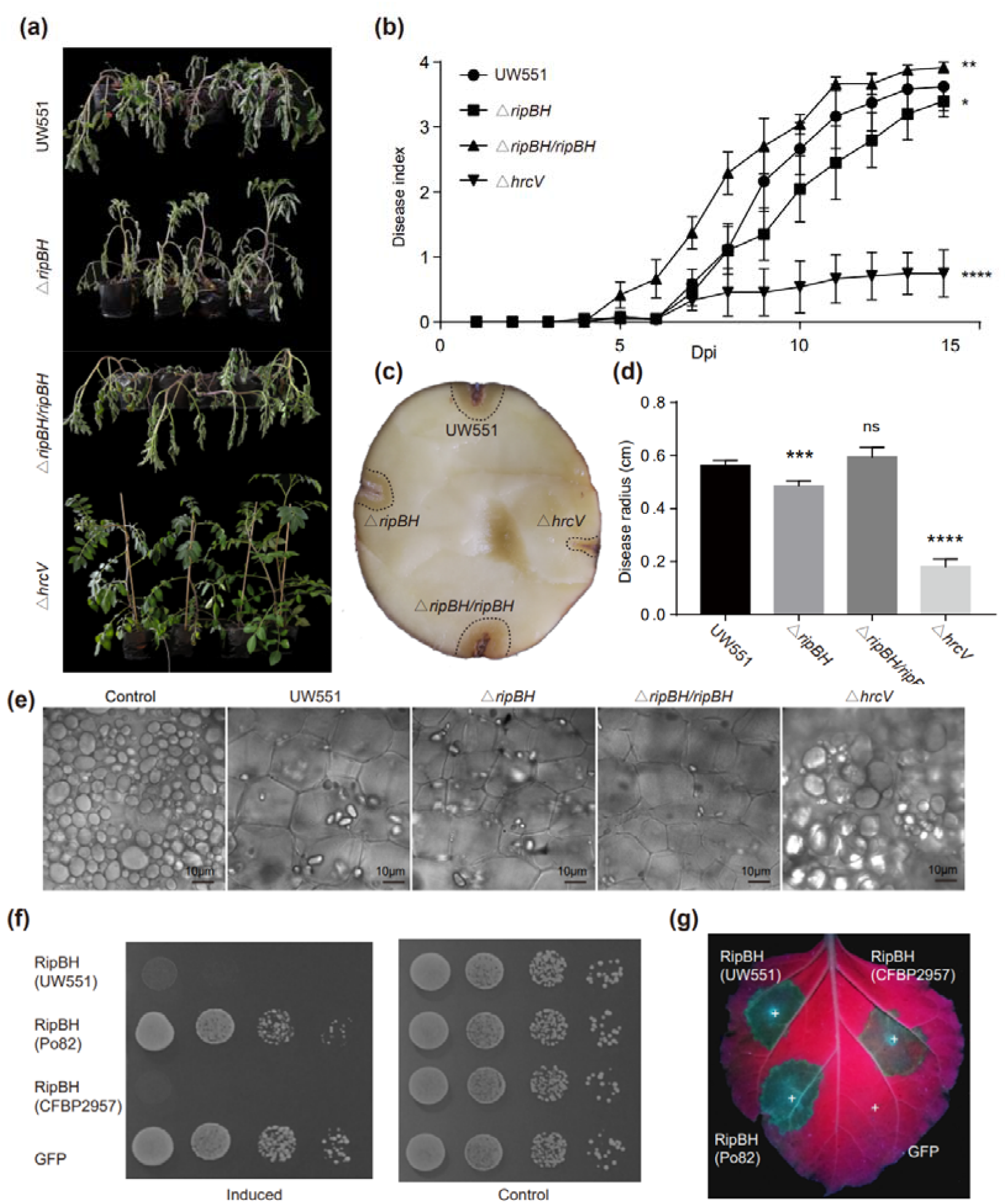
RipBH promotes tuber cell death caused by brown rot. (a-d) Soil-drenching inoculation assays in potato using *R. solanacearum* UW551, a ripBH mutant (Δ*ripBH*), and a *ripBH* complementation strain (Δ*ripBH/ripBH*). Representative photographs showing the disease symptoms of potato plants (a) and tubers (c) infected by strains *hrcV, ripBH*, Δ*ripBH:ripBH*, and UW551. The potato root-drenched infections were performed to evaluate the virulence of effector mutants. The images shown were obtained at 14 days post-inoculation (dpi). In (b) the results are represented as disease progression, showing the average wilting symptoms in a scale from 0 to 4. Values from 3 independent biological repeats were pooled together (mean ± SEM; n = 20; *, p < 0.05; **, p < 0.01; ****, p < 0.0001; Two-way ANOVA test). (d) The average radius of disease area in potato tubers were showed. Values from 3 independent biological repeats were pooled together (mean ± SEM; n = 15). Asterisks indicate significant differences (***, p < 0.001; ****, p < 0.0001; Student’s t-test). (e) The morphology of potato tuber after pathogen infection. Tuber section tissue near the inoculated site was visualized by microscopy. (f) Yeast growth inhibition assay showing serial dilutions of *S. cerevisiae* BY4743 cells grown under inducing (galactose) or control (glucose) conditions that are carrying plasmids expressing the *ripBH* homologous genes under the control of the GAL1 promoter. The cells were grown at 26 °C for 2 days for SD (−Ura) and 3 days for SGal (−Ura). (g) Transient expression assay of the *ripBH* homologous genes in *N. benthamiana*. The images were obtained at 72h after infiltration.

The induced expression system in yeast and the transient expression system in tobacco are two eukaryotic expression systems commonly used for bacterial effector phenotype screening (Fujiwara et al., 2016). Thus, we test the RipBH and two other homologs in these two systems. The results showed that RipBH (UW551) and RipBH (CFBP2957) could cause yeast growth inhibition. All three homologs could trigger cell death in *N. benthamiana* (Figure 1f, g). Unlike the hypersensitive response, RipBH-triggered cell death was not blocked by silencing the immune genes (SAG101, PAD4, NDR1) (Figure S2a, b), implying that the cell death triggered by RipBH was not a HR response. This data strongly suggested that RipBH showed strong cellular virulence in the yeast and tobacco cells.

To explore the biochemical functions of RipBH, we performed sequence analysis. The sequence alignments showed RipBH shares an identical N-terminal enzymic domain with EspL, a cysteine protease effector in the enteropathogenic *Escherichia coli* (EPEC) (Pearson et al., 2017). The sequence identity between RipBH and EspL is low (26%), but the secondary structure of the two proteins is almost identical (Figure 2a). They shared the same enzymic domain in N-terminal and ankyrin domain in C-terminal. Besides, the protein tertiary structure prediction showed the N-terminal enzymatic domain of RipBH is similar to the CA clan of papain-like cysteine protease AvrpphB (Zhu et al., 2004). The alpha fold prediction showed that the C-terminal of RipBH harbors a large ankyrin domain containing 10 ankyrin repeats, which is much larger in size than AvrpphB (Figure 2b). Although RipBH and EspL share a low sequence identity, they contain the same conserved catalytic active sites (C135S, D268A, H244A, N117A) (Figure 2a), implying that RipBH is a cysteine protease.

**Figure 2.**
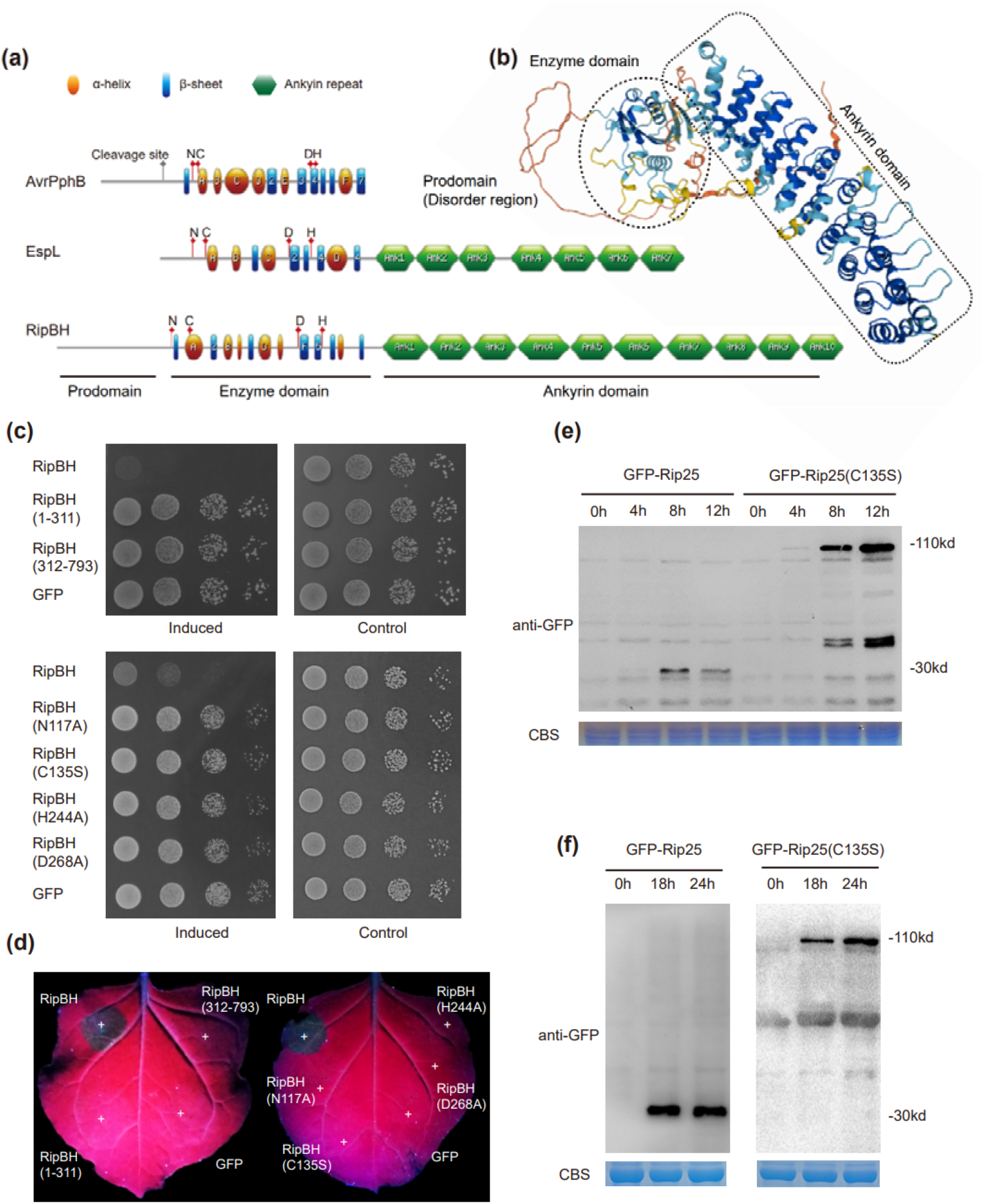
RipBH is an ankyrin cysteine protease. (a) Schematic representation of secondary structure of AvrpphB, EspL, and RipBH. (b) Structure predicted by AlphaFold. Color indicates a per-residue confidence score (pLDDT) between 0 (red) and 100 (blue). Three domains (prodomain, enzyme domain, and ankyrin domain) were shown. (c) Yeast growth inhibition assay showing serial dilutions of *S. cerevisiae* BY4743 cells grown under inducing (galactose) or control (glucose) conditions that are carrying plasmids expressing the *ripBH* genes truncated or mutated under the control of the GAL1 promoter. The cells were grown at 26 °C for 2 days for SD (−Ura) and 3 days for SGal (−Ura). (f) Transient expression assay of the truncated or mutated ripBH in *N. benthamiana*. The images shown were obtained at 72h after infiltration. (e and f) Immunoblot of protein extracts from induced expressed yeast (e) or agroinfiltrated leaves (f) of GFP-RipBH and GFP-RipBH(C135S). The protein loading is indicated by Coomassie Brilliant Blue (CBB) staining.

To study the biochemical function of RipBH, we mutated the catalytic active sites (C135S, D268A, H244A, N117A) of the N-terminal and generated truncated enzyme domain or ankyrin domain of RipBH. We found that the truncation of the entire enzyme domain or ankyrin domain terminal of RipBH abolished RipBH virulence function in yeast and tobacco (Figure 2c, d). Moreover, mutation of any catalytic active sites (C135S, D268A, H244A, N117A) abolished RipBH virulence function in yeast and tobacco as well (Figure 2c, d). This demonstrated that the N-terminal catalytic active site is indispensable for virulence.

In most cysteine proteases, the N-terminal prodomain usually acts as an endogenous inhibitor of the mature enzyme (Otto and Schirmeister, 1997). To activate the mature enzyme, removal of the prodomain is always necessary. This process is termed as auto-cleavage. We also detected the prodomain fragment (30kd) in immunoblot during GFP-RipBH expression in both yeast and tobacco (Figure 2e, f), implying that the prodomain of RipBH is cleaved during expression in the eukaryotic host cell. Moreover, this auto-cleavage is dependent on catalytic active sites. Mutation of catalytic active center (C135) abolished the auto-cleavage activities of GFP-RipBH (Figure 2e, f). Together, all these data strongly suggested that RipBH is a cysteine protease.

We investigated whether the large number (10) of ankyrin repeats in RipBH is critical for its functions. We generated a total of nine truncated RipBH proteins with different numbers of ankyrin repeats. The results showed all the truncation of ankyrin repeats compromised the ability to trigger cell death (Figure 3a, b). Except for RipBH (A1-5) with 5 ankyrin repeats showed a relatively weak cell death phenotype, all other truncation forms were unable to trigger cell death, suggesting that each ankyrin repeat is required for cell virulence (Figure 3a, b). We further analyzed the sequence of RipBH family members in other sequenced RSSC and found that most of the RipBH family members contain 10 ankyrin repeats, and only two family members contain 7 ankyrin repeats, suggesting that the number of ankyrin repeats are conserved and vital to virulence function (Figure S4).

**Figure 3.**
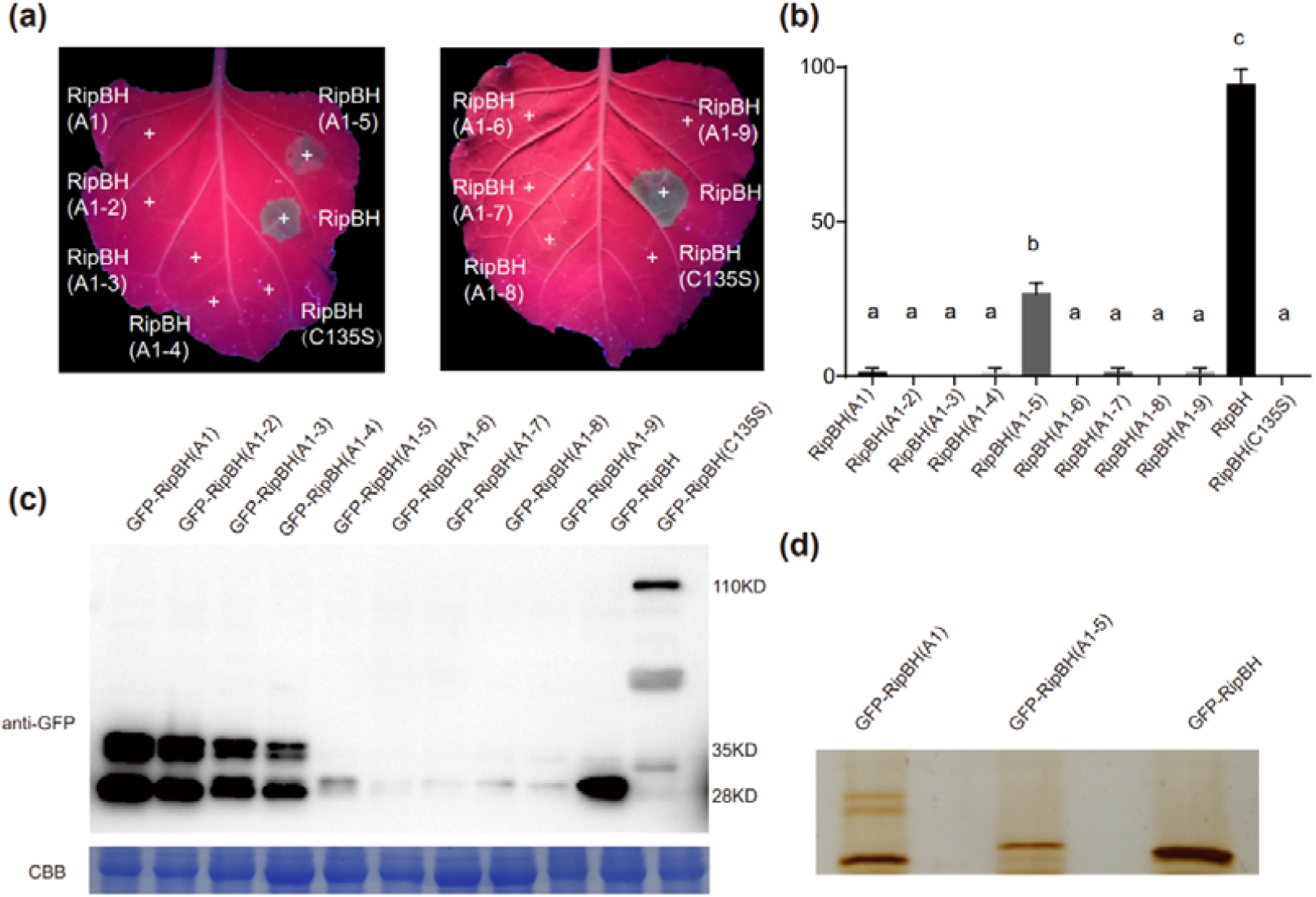
Ankyrin repeats is involved in the recognition of the cleavage site. (a and b) Transient expression assay of the ankyrin repeat truncated ripBH in *N. benthamiana*. The images shown were obtained at 72h after infiltration (a). (b) The barplot showed the cell death percentage of the transient expression site. Letters indicate significant differences (n=30, *p* < 0.01, One-way ANOVA test and Tukey’s multiple comparisons test). (c) Immunoblot of protein extracts from agroinfiltrated leaves of the ankyrin repeat truncated ripBH. The protein loading is indicated by CBB staining. (d) Silver staining of immunoprecipitated prodomain fragments of the ankyrin repeat truncated ripBH.

To test whether the ankyrin repeats are required for auto-cleavage, we also analyzed the auto-cleavage pattern of all the RipBH truncation proteins. During the expression of GFP-RipBH, only one protein fragment was detected by the immune blot assay (Figure 3c). Interestingly, we found three protein fragments appeared during the expression of a truncated protein with less than five ankyrin repeats including RipBH(A1), RipBH(A1-2), RipBH(A1-3), and RipBH(A1-4), which increase two more cleavage sites (Figure 3c). This implied the first 5 ankyrin repeats are involved in the recognition of the cleavage site. However, the other five truncated proteins RipBH(A1-5), RipBH(A1-6), RipBH(A1-7), RipBH(A1-8), and RipBH(A1-9) showed a relatively weak auto-cleavage activity (Figure 3c). We immunoprecipitated the N-terminal fragments and found the auto-cleavage pattern of a truncated protein with more than five ankyrin repeats is also slightly different from RipBH with full-length (Figure 3d). Together, these data strongly support that ankyrin repeats is involved in the recognition of the cleavage site and is indispensable for cell virulence.

Ankyrin repeat domain has a high binding affinity with proteins, nucleotides, and even membranes (Mosavi et al., 2004). Thus, we monitored the subcellular localization of GFP-RipBH (C135S) using a transient expression system in *N. benthamiana*. We found GFP-RipBH(C135S) widely distributed in the cytoplasm and nucleus (Figure 4a). Interestingly, the expression of the active form GFP-RipBH showed massive membrane bubbles in the cytoplasm (Figure 4a). We also found the same membrane bubbles during the expression of two homologous proteins (RipBH-PO82 and RipBH-IPO1609). To investigate whether it was associated with autophagosome, we co-expressed the autophagic marker GFP-ATG8i with RipBH. Interestingly, we observed plethora of autophagosomes during the co-expression with RipBH, but not with catalytic center mutant RipBH(C135S) (Figure 4b). The immunoblots against GFP also showed a strong autophagic flux that occurred during co-expression with RipBH instead of with RipBH(C135S) (Figure 4c).

**Figure 4.**
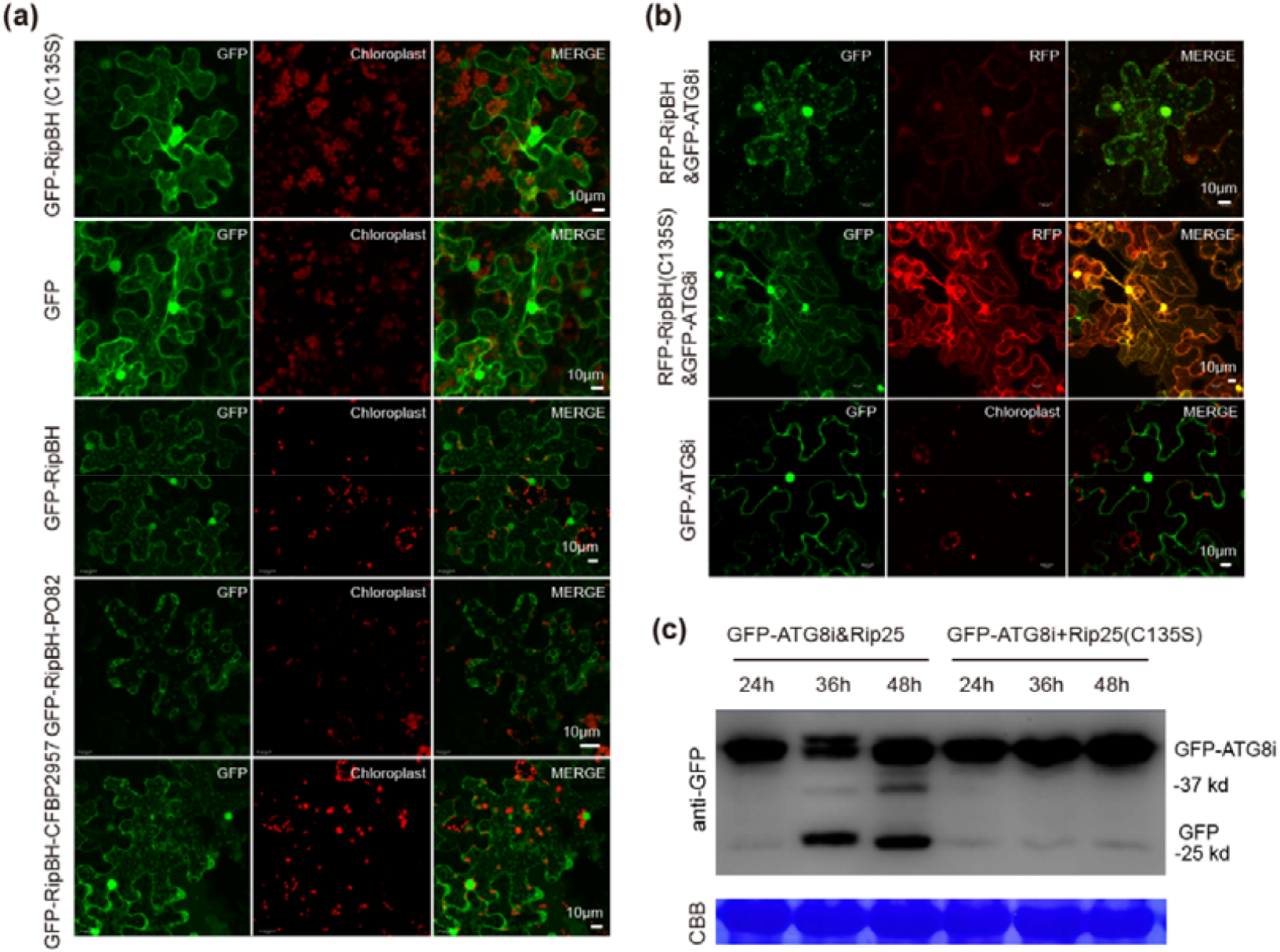
RipBH triggered the autophagy-associated cell death. (a) RipBH localized in the cytoplasm. The confocal images were captured at 36 h after transient expression in six-leaf-stage *N. benthamiana*. The white scale bar indicates 10 μm. (b) RipBH facilitated autophagosome formation. The confocal images were captured at 36 h after transient expression in six-leaf-stage *N. benthamiana*. The white scale bar indicates 10 μm. (c) Immunoblot showed the co-expression of RipBH and ATG8i in transient expression leaves. The protein loading is indicated by CBB staining.

To explore the relationship between autophagy and cell death triggered by RipBH, we silenced the autophagy-related genes *ATG5* and *ATG18a* to abolish the autophagy pathway and performed the cell death assay in these silenced tobaccos (Figure S5a). The results showed that blocking autophagy pathway decreased the cell death percentage significantly (Figure S5b), but not prevent the cell death triggered by RipBH completely. There are at least three types of autophagy-related cell death processes including autophagy-associated cell death and autophagy-dependent cell death (Denton and Kumar, 2019). The criteria to distinguish autophagy-associated cell death from autophagy-dependent cell death is that the blocking of autophagy, through either genetic or chemical means, prevents cell death completely (Denton and Kumar, 2019). In our study, we observed the autophagosome formation during the expression of RipBH, but silencing of the autophagy pathway was unable to stop the cell death process completely, suggesting the RipBH-triggered cell death is autophagy-associated cell death, not the autophagy-dependent cell death.

Potato brown rot, one of the most destructive diseases of potatoes, has been reported to affect about 3.75 million acres all over the world (Wang et al., 2017). Due to successive infection strategy, RSSC could enter potato root through the wound, colonizes the xylem, and even hide in the tuber causing wider spread. After potato harvest, RSSC is still capable to replicate in the tuber and causes cell death and necrosis eventually resulting in rotting tubers. However, few mechanisms of pathogenesis in tubers caused by brown rot were reported. Here, we first reported that a cysteine protease effector RipBH is required for tuber pathogenesis. RipBH is a novel structure cysteine protease with an ankyrin repeat domain and displays strong cell toxicity in yeast and tobacco. Interestingly, the ankyrin repeat truncation proteins still exhibited auto-cleavage activity but showed a different cleavage pattern, indicating the ankyrin repeat domain is not required for cleavage activity but is essential for cleavage site recognition. Although truncation proteins still exhibited auto-cleavage activity, yet reduced the cell toxicity in *N*.*benthamiana*, indicating that the ankyrin repeats play an essential role in the recognition of the target. Finding the recognition pattern would provide us with a potential target degradation tool. Unfortunately, we fail to find the target of this cysteine protease due to the poor solubility and the high toxicity. Exploring substrates of enzymes is a tough process, especially for the protease. Thus, in our further study we need to use other strategies to further explore the substrates of RipBH.

## ACKNOWLEDGEMENTS

We are grateful for the useful advice on this project from Alberto P. Macho and Shunping Yan. We thank Rui Jiang, Juan Du, and Qin He for fruitful discussion, especially for Qin He. We thank Caitilyn Allen and Philippe Prior for providing us with *R. solanacearum* strains. We also thank all members of the potato group in Huazhong Agricultural University for supporting this project. This work was funded by the National Natural Science Foundation of China (31871686) and the China Agriculture Research System of MOF and MARA (CARS-09-P07). No conflict of interest exits in the submission of this manuscript, and all authors approve the manuscript for publication.

## Suppleymentary figures

**Figure S1.**
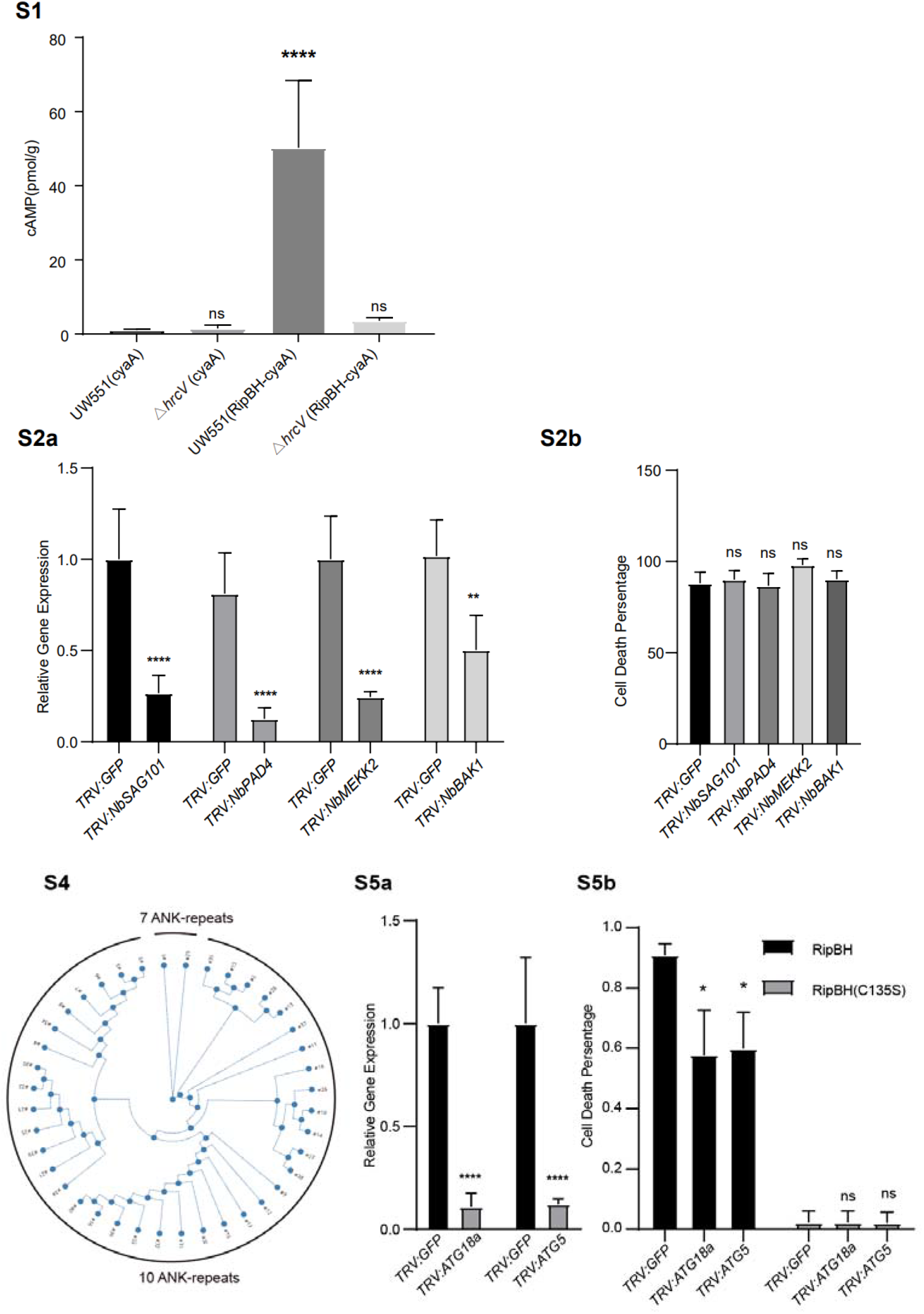
The RipBH is a type III-secreted effector. The cyclic adenosine monophosphate (cAMP) levels were measured at 7 days post-inoculation (n=4, ****, *p* < 0.001; Student’s t-test).

Figure S2 The ankyrin repeat number of RipBH homologs were conserved.

Figure S3 RipBH-triggered cell death was not blocked by silencing the immune genes. (a) Relative gene expression in plants subjected to virus-induced gene silencing. Plant leaves were sampled at 3 weeks after VIGS (n=3; ****, *p* < 0.001; **, *p* < 0.01; Student’s T-test). (b) The barplot showed the cell death percentage of the transient expression site. Asterisks indicate significant differences (n=30; ns, no significance; Student’s T-test).

Figure S4 The ankyrin repeat number of RipBH homologs were conserved.

Figure S5 Blocking autophagy pathway decreased the cell death percentage. (a) Relative gene expression in plants subjected to virus-induced gene silencing of *ATG18a, ATG5*. Plant leaves were sampled at 3 weeks after VIGS (student-t test, ****, *p* < 0.001, **, *p* < 0.01). (b) The barplot showed the cell death percentage of the transient expression site. Asterisks indicate significant differences (n=30; **, *p* < 0.05; ns, no significance; Student’s T-test).

