## Supplementary figures and images for "A toxic ankyrin cysteine protease effector RipBH of brown rot triggered autophagy-associated cell death"

### Supplemental figures

**S1**

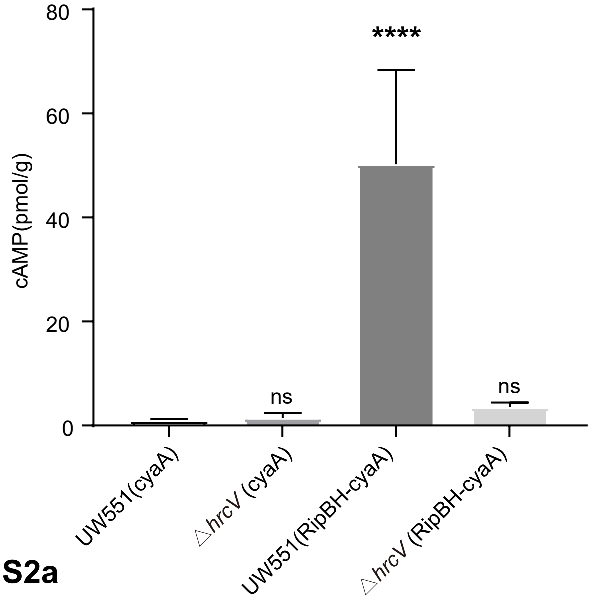


**S2a**

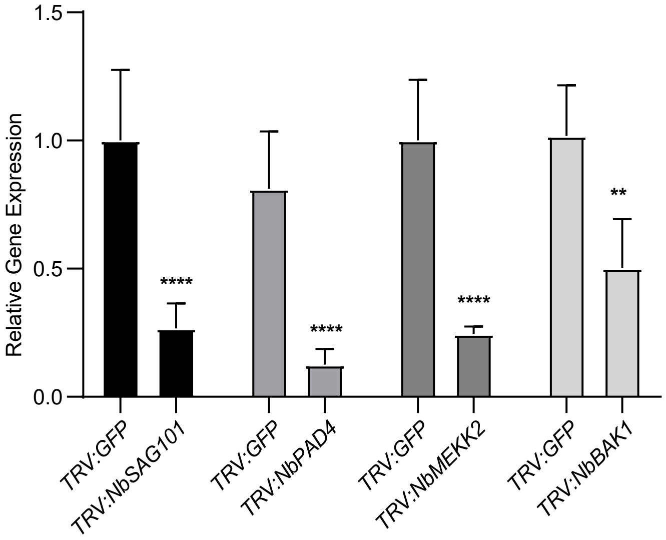


**S2b**

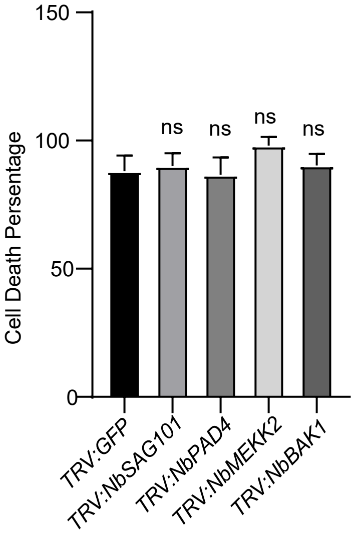


**S3**

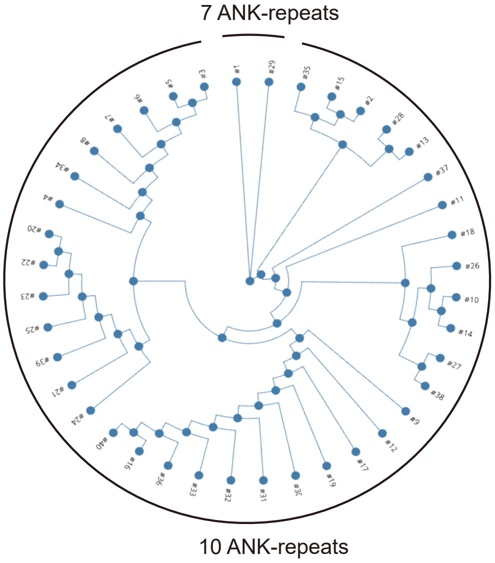


**S4a**

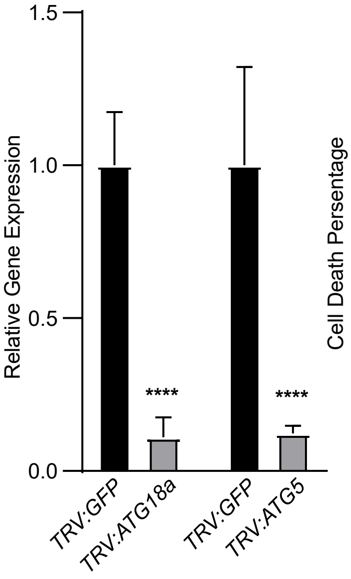


**S4b**

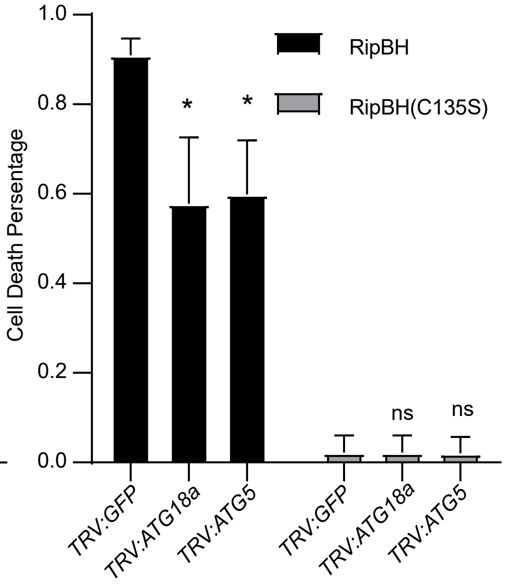
